# Engineered Genetic Redundancy Relaxes Selective Constraints upon Endogenous Genes in Viral RNA Genomes

**DOI:** 10.1101/323329

**Authors:** Silvia Ambrós, Francisca De la Iglesia, Sttefany M. Rosario, Anamarija Butković, Santiago F. Elena

## Abstract

Genetic redundancy, understood as the functional overlap of different genes, is a double-edge sword. At the one side, it is thought to serve as a robustness mechanism that buffers the deleterious effect of mutations hitting one of the redundant copies, thus resulting in pseudogenization. At the other side, it is considered as a source of genetic and functional innovation. In any case, genetically redundant genes are expected to show an acceleration in the rate of molecular evolution. Here we tackle the role of genetic redundancy in viral RNA genomes. To this end, we have evaluated the rates of compensatory evolution for deleterious mutations affecting an essential function, the suppression of RNA silencing plant defense, of tobacco etch potyvirus (TEV). TEV genotypes containing deleterious mutations in presence/absence of engineered genetic redundancy were evolved and the pattern of fitness and virulence recovery evaluated. Genetically redundant genotypes suffered less from the effect of deleterious mutations and showed relatively minor changes in fitness and virulence. By contrast, non-genetically redundant genotypes had very low fitness and virulence at the beginning of the evolution experiment that were fully recovered by the end. At the molecular level, the outcome depended on the combination of the actual mutations being compensated and the presence/absence of genetic redundancy. Reversions to wild-type alleles were the norm in the non-redundant genotypes while redundant ones either did not fix any mutation at all or showed a higher nonsynonymous mutational load.

## Introduction

Viruses are well known vectors of horizontal gene transfer (HGT) among hosts species (Penadés et al. 2015). In some well documented cases, viral genes have been inserted into cellular genomes, serving as sources of genetic and functional innovations (Koonin 2010; Liu et al. 2010). One of the most illustrative examples of such virus-mediated innovations is the acquisition of syncytin, an essential protein for the morphogenesis of placenta, from an endogenous retrovirus at the origin of the lineage of placental mammals (Mi et al. 2000). Gene trafficking goes in both directions, and viruses sometimes acquire genes from the genome of their host cells, or from other viruses coinfecting the same host. For example, phylogenetic studies have suggested that many *Mimivirus* genes have been acquired by HGT from the host (Moreira and Brochier-Armanet 2008). A second good example of HGT from host to virus is the homologue of the 70 kDa heat shock protein encoded in the genome of plant closteroviruses (Dolja et al. 2006). Although HGT is thought to be an essential mechanism for virus evolution, strong selection against increasing genome size may hinder its importance, especially for small RNA viruses (Zwart et al. 2014, Willemsen et al. 2016b). Accommodation of a new gene into the already compacted genome of the recipient virus would necessarily involve changes in the regulation of gene expression, re-tuning interactions among all viral proteins and between them and the host proteome, and optimization of codon usage of the newly acquired gene to the characteristic bias of the recipient virus. Right upon insertion, the exogenous sequence will generally have a negative impact in the fitness of the recipient virus (Dolja et al. 1993; Zwart et al. 2014). Despite any possible long-term possible benefit, shortsighted selection will purge these low fitness genomes from a population numerically dominated by the wild-type virus. Even in isolation, these longer genomes will have a disadvantage relative to deletion mutants that will be generated during their replication (Zwart et al. 2014; Willemsen et al. 2016b).

In a recent series of studies, we have explored the constraints to innovation in the prototypical genome of a plant (+)-sense ssRNA monopartite virus belonging to the picorna-like superfamily, *Tobacco etch virus* (TEV; genus *Potyvirus*, family *Potyviridae*). More precisely, we have explored the short-term cost of acquiring new exogenous genes and their evolutionary fate (Zwart et al. 2014; Willemsen et al. 2016b; Willemsen et al. 2017), the fitness burden imposed by changes in gene order (Majer et al. 2014; Willemsen et al. 2016a) and the effect on virus fitness of the HGT of an essential viral gene into the host genome and its constitutive expression (Tromas et al. 2014). In addition, using a segmented plant RNA virus, *Alfalfa mosaic virus*, we have also been exploring the fitness consequences of genome segmentation and multipartition (Sánchez-Navarro et al. 2013; Wu et al. 2017). For its direct relationship with the work reported in this article, we would mention here the cases in which we added new genes into TEV genome. We generated a number of artificial genomes carrying insertions of heterologous sequences from different origins. First, the *AlkB* domain from *Nicotiana tabacum*, which is involved in correcting alkylation damages in nucleic acids (Willemsen et al. 2017). Second, the *2b* gene from *Cucumber mosaic virus* (CMV), which is a suppressor of RNA silencing (RSS) (Willemsen et al. 2017). Third, the *eGFP* gene encoding for a version of the green fluorescent protein (Zwart et al. 2014). The first two cases would provide the virus with completely new *(AlkB)* or redundant (2b) functions, whereas the *eGFP* would, in principle, not provide any useful function to the virus but just increase its genome length. In all cases, engineered genomes were evolved by serial passages in *N. tabacum*, with at least five independent evolutionary lineages. At the end of the experimental evolution, the evolved viral populations were phenotypically (viral load, relative fitness, infectivity, and virulence) and genetically (Illumina NGS study of genetic variability within each evolved lineage) characterized. In summary, *eGFP* and *AlkB* genes were always removed from the genome at a rate that depended on the demographic conditions, recombination rate, cloning site, and the exogenous gene itself (Willemsen et al. 2018). Interestingly, the *2b* RSS was pervasively retained in all evolved lineages (Willemsen et al. 2017). Furthermore, functional analyses demonstrated that the 2b protein was able of complementing the effect of strong and weak hyposuppressor mutants negatively affecting the TEV RSS multifunctional protein HC-Pro (Willemsen et al. 2017). Therefore, the engineered TEV/2b virus encodes for evolutionarily stable genetic redundancy in its genome. In this context, *sic* “genetic redundancy means that two or more genes are performing the same function and that inactivation of one of these genes has little or no effect on the biological phenotype” (Nowak et al. 1997). Genetic redundancy may open the possibility, at least in theory, for the HC-Pro multifunctional protein to relax purifying selection on protein domains involved in the RSS function and to optimize some of its other activities as protease, cell-to-cell movement or as helper-component during aphid horizontal transmission (a trait that should be irrelevant in our experimental system, because transmission is mechanical) (for a recent review on HC-Pro functions, see Valli et al. 2018).

To experimentally explore the role of genetic redundancy, provided by *2b*, on the evolutionary fate of a strong (hereafter named as FINK; see below) and a weak hyposuppressor (hereafter named as AS13; see below) mutants affecting the RSS activity of HC-Pro, we have performed an evolution experiment starting with two different TEV genetic backgrounds. The first consists on the wild-type TEV genotype into which we introduced the FINK and AS13 mutations, respectively. The second genetic background consists on the TEV/2b engineered genotype into which we introduced the same set of mutations. We sought to test the following hypotheses: (*i*) genetic redundancy will compensate for the inefficient RSS activity of HC-Pro hyposuppressor mutants, slowing down the rate of compensatory (phenotypic and genotypic) evolution compared with the non-redundant case. (*ii*) By buffering the effect of mutations in HC-Pro RSS activity, mutational load in this sequence may further increase.

## Materials and Methods

### Viral Constructs and Stocks

Mutant genotypes TEV-FINK, TEV-FINK/2b, TEV-AS13, and TEV-AS13/2b were constructed by site-directed mutagenesis as previously described (Torres-Barceló et al. 2008; Willemsen et al. 2017). TEV/2b virus variants contain the *2b* gene from CMV. FINK and AS13 mutant genotypes contain nucleotide substitutions within the central RSS region of the HC-Pro protein. The functional strong hyposuppressor mutant FINK consists in the G548U nucleotide substitution leading to the R183I amino acid change (positively-charged hydrophilic by non-charged hydrophobic) in the highly conserved FRNK motif within the RNA-binding domain (RBD) A (Shiboleth et al. 2007; Kung et al. 2014). The functional weak hyposuppressor AS13 double mutant contains the A896C and A899C nucleotide substitutions leading to E299A and D300A amino acid changes (negatively-charged and hydrophilic by non-charged hydrophobic) within the RBD B domain (Kasschau et al. 1997). The viability and fitness effects of these mutants, relative to the wild-type TEV, were assessed in detail in previous works (Torres-Barceló et al. 2008; Willemsen et al. 2017). Homogenized stock tissue from plants infected with these four viral genotypes was preserved at −80 °C and used as stock for initiating the evolution experiments described below.

### Experimental Evolution

A variable number of independent evolution lineages (as indicated in parentheses) were stablished for each of the five engineered genotypes: TEV-FINK (2), TEV-FINK/2b (5), TEV-AS13 (3), and TEV-AS13/2b (5). In all cases, 100 mg of the corresponding ancestral tissue-infected stocks were resuspended with an equal volume of phosphate buffer (50 mM KH_2_PO_4_, pH 7.0, 3% polyethylene glycol 6000) and 5 μl of this mixture used to mechanically inoculate the third true leaf of 20 4 or 5-week-old *N. tabacum* L. cv Xanthi *NN* plants dusted with Carborundum. In the case of TEV-AS13, no infected plants were obtained for one lineage and thus we used the other successful lineage (L2) to create a replicate (L3) that was maintained independently afterwards. Inoculated plants were kept in a Biosafety Level-2 greenhouse chamber at 25 °C under a 16 h natural sunlight (supplemented with 400 W high-pressure sodium lamps as needed to ensure a minimum light intensity of PAR 50 μmol/m^2^/s) and 8 h dark photoperiod. The passage duration was determined by the appearance of at least one symptomatic plant per lineage between 9 and 15 days-post infection (dpi). Those lineages in which no plant displayed symptoms 15 dpi, were left to grow for five extra days. Leaves above the inoculated leaf were collected from all symptomatic plants, stored at −80 °C and used for subsequent serial passages. Then, 100 mg of plant tissues were homogenized into a fine powder with liquid N2 using a Mixer Mill MM400 (Retsch GmbH, Haan, Germany). At each passage, the third true leaf of 10 (for TEV-FINK/2b and TEV-AS13/2b) or 15 - 20 (for TEV-FINK and TEV-AS13) 4 or 5-week-old plants were mechanically inoculated as just described. For those lineages in which the viral load was very low, volumes of phosphate buffer were reduced in order to equilibrate the genome equivalents of all evolved viral lineages. A total of five serial passages were conducted.

### Evaluation of Virulence Traits

After inoculation, plants were observed every day for the presence of symptoms and the number of symptomatic plants recorded. Three different virulence-related traits were estimated from these data. First, the percentage of symptomatic plants at the end of the experiment (*t* = 20 dpi) was used as an estimate of infectivity (*i*). Second, infectivity time-series data were submitted to a Kaplan-Meier regression analysis of survival times and the median time to the appearance of symptoms (ST_50_) estimated. Third, using the infectivity time-series data, we also estimated the area under the disease progress stairs curve *(AUDPS)* (Simko and Piepho 2012):

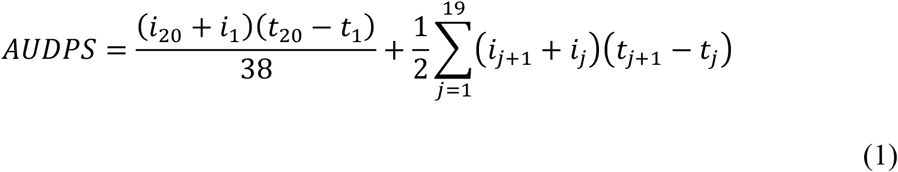

where *i*_*k*_ stands for the frequency of infected plants at *k* dpi.

Fourth, each day the severity of symptoms (*S*) was recorded for each individual plant according to the following semi-quantitative scale: *S* = 0, no symptoms; *S* = 1, curly leaves and disperse chlorotic spots; *S* = 2, incipient vein etching; *S* = 3, extensive and strong etching and necrotic tissue; and *S* = 4, generalized necrosis and plant death. The average severity of symptoms, *S̄*, as evaluated at each dpi (*t*) for the set of plants was fitted to the following logistic regression model (Campbell and Madden 1990):

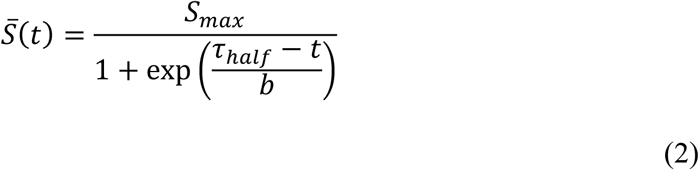

where *S*_*max*_ represents the maximal symptoms score value, *τ*_*half*_ the half-time elapsed from inoculation to observing the maximal symptoms developed and *b* stands for the steepness of the symptoms development curve, *i.e.*, the inverse of the speed at which plants develop the maximal symptoms.

The six virulence-related traits evaluated for each lineage, *L*, after each evolution passage, *P*, were analyzed using multivariate analysis of variance methods (MANOVA) and fitted to the following multivariate linear model:

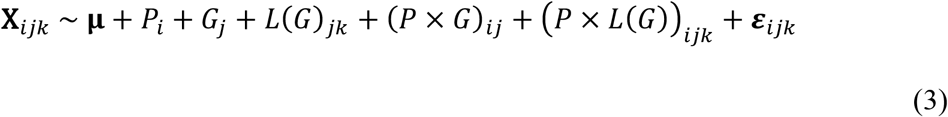

where **X**_*ijk*_ is the vector of observed trait values for lineage *k* of viral genotype *j* observed at passage *i*, **μ** the vector of grand mean values, ***ε***_*ijk*_ the vector of errors assumed to be distributed as a multivariate Gaussian. *G* is the factor accounting for the virus genotype (*i.e.*, TEV-FINK, TEV-FINK/2b, TEV-AS13 and TEV-AS13/2b), *L* is nested within *G* [*L*(*G*)] and *P* was considered as a covariable. Interactions between the covariable and the two factors have also been incorporated into the model. The significance of each component in the model has been evaluated with the Wilk’s *∧* statistic. Associations between virulence traits were evaluated using a partial correlation coefficient controlling for *G*, *r*_*p*_. A factorial analysis was also performed and the six variables were reduced into a single principal component (*PC1*) that explained up to 79.73% of total variance. For the shake of summarizing the evolution of virulence into visually appealing plots, *PC1* will be presented instead of each single trait.

All statistical analyses described in this section and hereafter were performed using SPSS version 23 software (IBM, Armonk NY, USA).

### Evaluation of Within-Host Fitness

Viral load was evaluated as a measure of within-host replicative fitness. Viral load was only determined after passages 1 and 5. Total plant RNA extractions (RNAt) were performed from 100 mg of tissue powder with the Plant RNA Isolation Mini kit (Agilent Technologies, Santa Clara CA, USA). Viral accumulation (TEV genomes per ng of RNAt) was quantified by RT-qPCR as previously described (Zwart et al. 2014). Briefly, reactions were performed in 20 μl volume using the GoTaq 1-Step RT-qPCR System (Promega, Fitchburg WI, USA) following the instructions recommended by the manufacturer in an ABI StepOne Plus Real-time PCR System (Applied Biosystems, Foster City CA, USA). The cycling conditions were as follow: an RT phase of 15 min at 37 °C followed by 10 min at 95 °C; a PCR phase consisting in 40 cycles of 10 s at 95 °C, 34 s at 60 °C and 30 s at 72 °C; and a final phase of 15 s at 95 °C, 1 min at 60 °C and 15 s at 95 °C. The following specific TEV primers designed in the *CP* region were used: forward 5’-TT GGTCTTGAT GGCAACGT G-3’ and reverse 5’-TGTGCCGTTCAGTGTCTTCCT-3’. The StepOne Software v.2.2.2 (Applied Biosystems) was used to analyze quantitative data. All ancestral and evolved viral lineages were normalized in order to equilibrate genome equivalents in the sample viral inoculums of the first and second passages.

Viral load data were fitted to a generalized linear model (GLM) with a Gamma distribution and a logarithmic link function (based on the smallest *BIC* value among a set of alternative models). The model equation is now written as

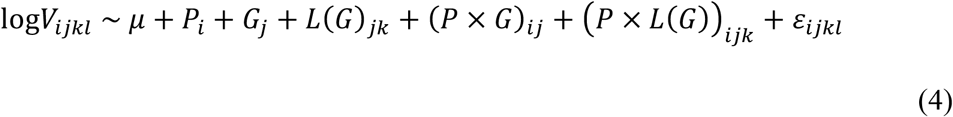

where *V*_*ijkl*_ stands for the viral load for the *l*-th replicate plant (3) from lineage *k* of genotype *j* at passage *i* (in this case 1 or 5) and all other factors are as described in the previous section. The significance of each component in the model was evaluated by means of likelihood ratio tests *(LRT)* that asymptotically follow a *χ*^2^ probability distribution. The magnitude of effects was evaluated using the 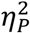 statistic (proportion of total variability in viral load attributable to each factor in the model). Conventionally, values 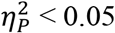 are considered as small, 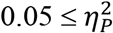 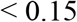 as medium and 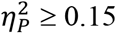 as large effects.

All raw data are available at LabArchives under doi: 10.6070/H418352M.

### Evaluation of Insert Stability by RT-PCR

Reverse transcription and amplification reactions (RT-PCR) were performed in 20 μl volume using the GoTaq 1-Step RT-qPCR System (Promega) as indicated before. Input RNAt samples consisted in 150 to 300 ng from the selected plants per each lineage of passages 1 and 5 and two primers flanking the *2b* gene region: forward PC96-97F 5’-TACGATATTCCAACGACTG-3’ and reverse PC92-96R 5’-GCAAACTGCTCATGTGTGG-3’. Amplification protocol was as follows: an RT phase of 15 min at 45 °C plus 2 min at 95 °C and a PCR phase consisting in 3 cycles of 15 s at 95 °C and 1 min at 67 °C and 37 cycles of 15 s at 95 °C and 1 min at 65 °C. Size and quality of PCR products were confirmed by electrophoresis on 1.5% agarose gels. TEV mutants rendered products of 651 pb or 298 pb according with the presence or absence of *2b*, respectively.

### Sequencing

To generate the consensus sequence of each mutant and lineage from passages 1 and 5, accurate RT-PCR products were obtained. RT reactions for the *HC-Pro* and *2b* genes were performed using the AccuScript Hi-Fi reverse transcriptase (Agilent Technologies) and reverse primers HCPro-R2545 5’-AATTGAAGGGGAGACTATTGCCAG-3’, that flanks the TEV *HC-Pro* cistron, or the PC92-96R flanking the 5’ *CP* cistron following manufacturer’s instructions. Then, viral genomes were amplified using Phusion DNA polymerase (Thermo Scientific, Waltham MA, USA), 1 μl of the RT reactions and primers PC96-97F and PC92-96R (*2b* gene) or HCPro-F 5’-GGAT GGGAT GTTGGT GGAT GCTC-3’ and HCPro-R 5’-CCTTGTGTGACCATATCTCGGTTC-3’ (*HC-Pro* gene). PCR protocol for *HC-Pro* and *2b* amplicons was as follows: initial denaturation 30 s at 98 °C, five cycles with denaturation step of 10 s at 98 °C, annealing step of 20 s at 57 °C and elongation step of 30 s at 72 °C followed with 27 cycles in the same conditions except that annealing step was undergone at 65 °C. Amplicon sizes, 1473 bp for *HC-Pro* and 651/298 for *2b*, were confirmed by electrophoresis on 1% agarose gels. Consensus sequences of all evolved lineages at passages 1 and 5 were obtained using Sanger sequencing and an ABI 3130 XL Genetic Analyzer (Applied Biosystems) in IBMCP’s DNA Sequencing Unit (www.ibmcp.csic.es/en/services/dna-sequencing-unit). Sequences were assembled using GENEIOUS version 11 software (www.geneious.com).

## Results

### Evolution of Virulence-related Traits

Fig. 1A illustrates some representative examples of the changes in symptomatology observed along the evolution experiment. For instance, panels b and c compare the drastic difference in the type of symptoms induced by TEV-AS13, consisting in disperse chlorotic spots (would correspond to *S* = 1 in the symptoms scoring system described in Materials and Methods section), and TEV-AS13/2b which shows a more generalized spotted chlorotic disease (*S* = 2) but without the characteristic etching of wild-type TEV (see plants on the right in panel g). Panel d shows the symptoms generated by lineage L4 of TEV-FINK at passage 3, again consisting in disperse chlorotic spots (*S* = 1), while lineage L1 of TEV-FINK at the same passage had already reverted to the typical etching and vein necrosis of TEV (panels e and f; *S* = 3). Panel g shows the differences between a non-infected plant (left; also panel a) and two plants infected with lineage L1 of TEV-FINK after the last passage (*S* = 3). In summary, both viral genotypes expressing the 2b RSS protein induced stronger symptoms than their corresponding counterparts. In all lineages, symptoms severity increased during the evolution experiment.

**Fig. 1.**
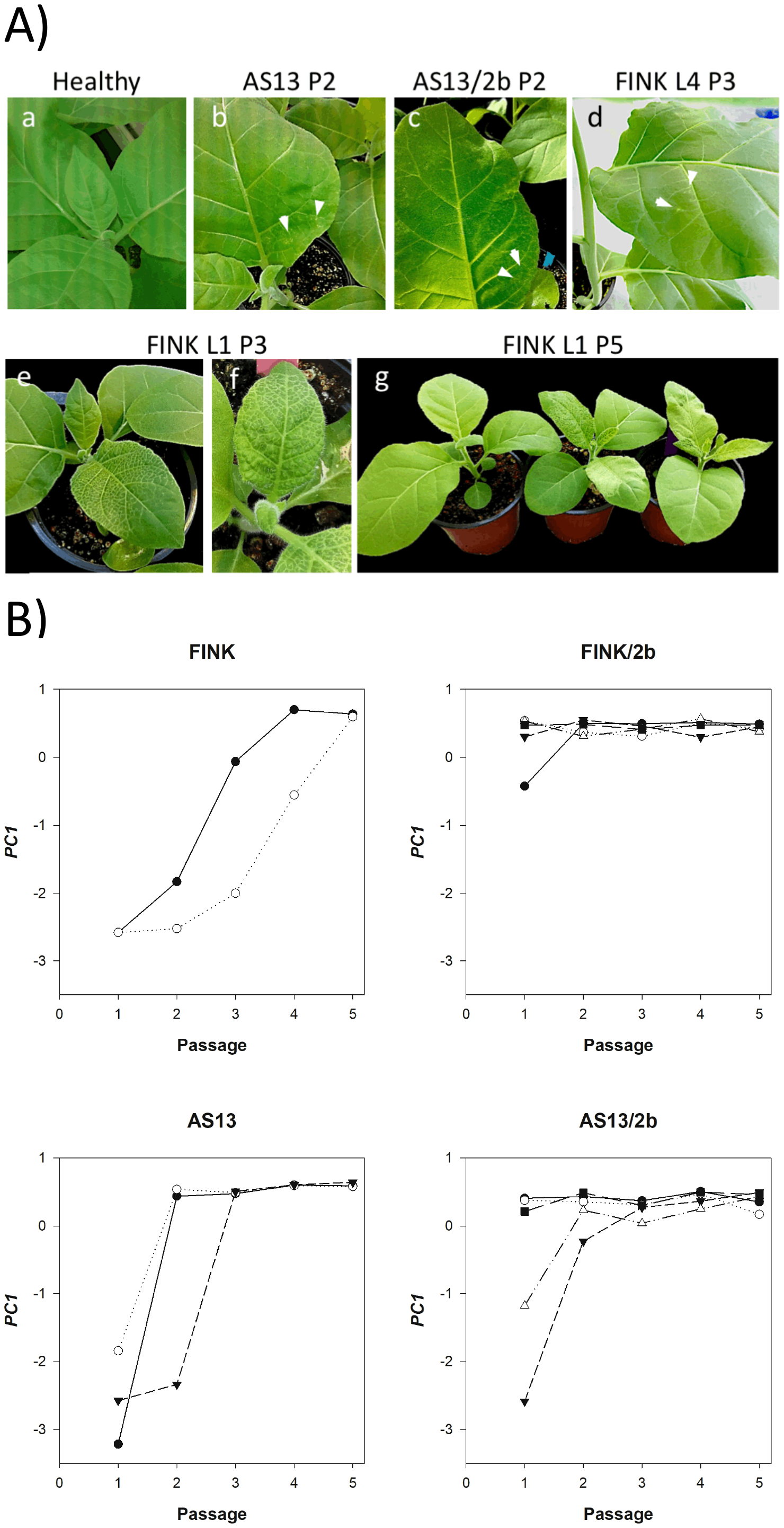
(A) Symptoms induced by the ancestral and evolved TEV genotypes. Upper panel: from left to right, (a) healthy plant, (b) chlorotic spots observed in a plant of passage 2 (P2) infected with one of the TEV-AS13 lineage, (c) the same for a plant infected with a one of the TEV-AS13/2b lineages, and (d) a plant from the third passage (P3) infected with lineage L4 of TEV-FINK. White arrows indicate the location of some chlorotic spots. Lower panel: (e) and (f) plants of passage P3 infected with lineage L1 of TEV-FINK showing the typical etching of TEV, (g) comparison of plants from the fifth passage (P5): healthy (left) or infected with lineage L1 of TEV-FINK (right), thus displaying no symptoms or the typical etch mosaic of TEV, respectively. (B) Evolution of the first principal component (*PC1*) summarizing the six virulence-related traits. See text for a detailed description of the contribution of the different traits into *PC1*. Different symbols correspond to the different independently evolved lineages.

After each passage, infectivity (*i*) and symptoms severity data (*S*) were fitted to equations (1) and (2), respectively; five different virulence-related traits were inferred: *i*, *AUDPS*, *S*_*max*_, *τ*_*half*_, and *b*. In addition, time-course infectivity data were also analyzed using the Kaplan-Meier regression method and the median time to infection, *ST*_*50*_, was also estimated. First, we explored the dependencies and associations between these six virulence-related traits using partial-correlation coefficients controlling for the viral genotype (*G*) factor. Table 1 shows the associations obtained. Partial correlations identify two groups of virulence-related traits that respond in a highly parallel manner. In the first group, *AUDPS*, *S*_*max*_ and *i* all show significant positive correlations. In the second group *ST*_*50*_, *τ*_*half*_ and *b* show significant pairwise positive correlations. However, correlations between traits belonging to different groups are all significantly negative (table 1).

**Table 1.** Partial correlations (*r*_*p*_), controlling for viral genotype, between the different virulence-related traits measured. In all cases 72 d.f. and *P* < 0.001.

|  | *i* | *AUDPS* | *ST*<sub>50</sub> | *S*<sub>max</sub> | *τ*<sub>half</sub> | *b* |
| --- | --- | --- | --- | --- | --- | --- |
| *i* | 1 | 0.841 | −0.841 | 0.847 | −0.424 | −0.645 |
| *AUDPS* |  | 1 | −0.997 | 0.989 | −0.506 | −0.834 |
| *ST*<sub>50</sub> |  |  | 1 | −0.989 | 0.502 | 0.835 |
| *S*<sub>max</sub> |  |  |  | 1 | −0.483 | −0.811 |
| *τ*<sub>half</sub> |  |  |  |  | 1 | 0.716 |
| *b* |  |  |  |  |  | 1 |

*i*: infectivity; *AUDPS*: area under the disease progress stairs curve; *ST*<sub>50</sub>: time to observe infections symptoms inferred from a Kaplan-Meier survival regression analysis; *S*<sub>max</sub>: maximal score value for the observed symptoms; *τ*<sub>half</sub>: half-time to develop the maximal symptoms; *b*: steepness of the symptoms development curve.

To explore the overall effect of different factors on these virulence-related traits, we fitted the MANOVA model shown in equation (3) to the multivariate data. Table 2 shows the results of this analysis. The covariable and the two factors in the linear model are all significant: virulence increases with passages, overall differences among viral genotypes exist and significant heterogeneity exist among lineages started from the same ancestral viral genotype (table 2). In addition, the interaction between passage (covariable) and genotype was also highly significant, indicating that the rates of virulence recovery were different between viral genotypes (table 2). Finally, within each viral genotype, lineages also differed in their rates of virulence evolution, although in this case the significance level is moderate (table 2; *P* = 0.021).

**Table 2.** Multivariate analysis of variance (MANOVA) of the six virulence-related traits. The model is described by equation 3.

| Source of variation | Wilk's $\Lambda$ | $F$ | d.f. hypothesis | d.f. error | $P$ |
| --- | --- | --- | --- | --- | --- |
| Intersection | 0.001 | 7669.225 | 6 | 40 | < 0.001 |
| Passage | 0.267 | 18.267 | 6 | 40 | < 0.001 |
| Genotype | 0.115 | 7.265 | 18 | 113.622 | < 0.001 |
| Lineage(Genotype) | 0.070 | 2.143 | 66 | 219.490 | < 0.001 |
| Passage×Genotype | 0.230 | 4.294 | 18 | 113.622 | < 0.001 |
| Passage×Lineage(Genotype) | 0.141 | 1.469 | 66 | 219.490 | 0.021 |

Given the strong dependencies among the six virulence-related variables, we applied a principal components analysis and found that 79.73% of the observed variability was explained by the major eigenvalue (*λ*_1_ = 4.784) of the variance-covariance matrix (the second eigenvalue was *λ*_2_ = 0.828 < 1 and negligible). Therefore, to represent the six variables into a single summary statistic, we computed the first principal component in the direction of *λ*, which results in the following expression:

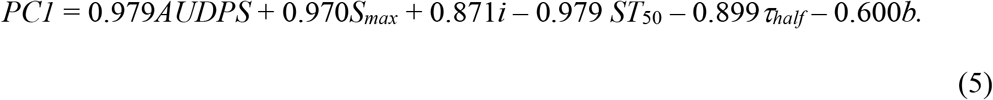

The biological meaning of this *PC1* is straightforward: it gives positive weights to *AUDPS*, *S*_*max*_ and *i* and negative weights to *ST*_*50*_, *τ*_*half*_ and *b*, in accordance with the results of the correlation analyses shown above. The three first variables measure frequency and severity of symptoms while last three are related with the time necessary for symptoms to be developed. In other words, evolving viruses are rising in virulence by increasing the positively weighted traits (*i.e.*, increasing infectivity or the severity of symptoms) and/or by reducing the negatively weighted ones (*i.e.*, shorter times to develop symptoms). *PC1* < 0 values would correspond to viruses that produce very few infected plants and of weak symptoms that take long to be observable. By contrast, *PC1* > 0 values will be characteristic of viruses that are highly infectious and quickly induce strong symptoms. Fig. 1B illustrates the evolution of virulence as described by *PC1:* a faster recovery of virulence in TEV-AS13 and TEV-FINK compared to their respective *2b*-carrying counterparts. However, after the first passage, a larger amount of variance exists among the lineages carrying the *2b* than among the non-carrying ones (fig. 1B). Lineage L1 of TEV-FINK/2b shows a lower *PC1* than the other four lineages TEV-FINK/2b; this difference was erased after passage 2. Likewise, lineages L8 and L11 of TEV-AS13/2b also show remarkable low values of *PC1* after the first passage, much lower than the other two TEV-AS13/2b lineages and in the same range of values observed for the genotypes not expressing the 2b RSS. Thereafter passage 3 differences among the five TEV-AS13/2b lineages disappeared.

To gain further insights into the role of RSS genetic redundancy during the evolution of virulence, we performed pairwise *post hoc* Bonferroni tests using *PC1* as variable in an ANOVA with the same structure than equation 3. Firstly, we compared the difference between TEV-FINK and TEV-FINK/2b (fig. 1B, panels FINK *vs* FINK/2b). The mean value of *PC1* for TEV-FINK was −1.018 ±0.101, while the mean value for TEV-FINK/2b was 0.411 ±0.064, being the difference highly significant (*P* < 0.001). Therefore, the expression of 2b buffers the strongly deleterious effect of the mutation R182I in the highly conserved FRNK motif of the RBD A domain of HC-Pro, which leads to a weak infection phenotype and hyposuppression activity in other potyviruses (Lin et al. 2007; Shiboleth et al. 2007; Kung et al. 2014), thus relaxing selection to revert or compensate this amino acid replacement. By contrast, in absence of a backup for the RSS activity, strong selection operates upon TEV-FINK to recover its ancestral HC-Pro activity which also correlates with a stronger symptom induction and virulence. Secondly, we compared the difference between TEV-AS13 and TEV-AS13/2b (fig. 1B, panels AS13 *vs* AS13/2b). The mean value of *PC1* for TEV-AS13 was −0.264 ±0.186, while the mean value for TEV-FINK/2b was 0.152 ±0.144, being this difference not significant (*P* = 0.092). This result suggests that 2b is not compensating the effect of the deleterious E299A/D300A mutations in the RBD B domain of HC-Pro to the same extent than it does for the R182I mutation in the RBD A domain, and thus selection may operate both in TEV-AS13 and TEV-AS13/2b to improve HC-Pro activity. Thirdly, we compared the difference between TEV-FINK and TEV-AS13 (fig. 1B, panels FINK *vs* AS13). In agreement with our above argumentation, these two genotypes show statistically similar profiles of virulence recovery (*P* = 0.050). Finally, we compared the differences between TEV-FINK/2b and TEV-AS13/2b (fig. 1B, panels FINK/2b *vs* AS13/2b). Supporting our above conclusion that genetic redundancy provided by 2b buffers deleterious mutational effects in the FRNK motif but not in the RBD B domain, the mean *PC1* values of these two genotypes are significantly different (*P* = 0.009), being greater for TEV-FINK/2b. In subsequent sections we will explore the molecular basis of this evolutionary recoveries.

### Compensatory Evolution of Within-Host Replicative Fitness

As a proxy to within-host viral fitness, we have quantified viral load by RT-qPCR (number of viral genomes per ng of RNAt extracted from 100 mg of plant tissue), as described in Materials and Methods, after the first and last evolution passages. The resulting values are shown in fig. 2A. In all cases, significant improvements in viral load has been observed. Table 3 shows the results of fitting the GLM model described by equation 4 to the data. The three factors have a highly significant effect on viral load. First, viral load at passage 5 is always greater than at passage 1 and the effect is of very large magnitude (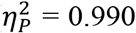). The largest differences are observed for the two lineages of TEV-FINK and lineage L5 of TEV-AS13 (fig. 2A). Second, consequently, the magnitude of improvement in viral load significantly varies among viral genotypes, being on average larger for TEV-FINK and TEV-AS13 than for the corresponding genotypes carrying the *2b*. Nonetheless, the magnitude of this effect is the smallest observed (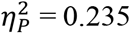) yet still can be considered as large. Third, heterogeneity can be observed between independent lineages evolved from the same ancestral genotype, being the effect also very large in magnitude. Fourth, the magnitude of these two effects actually depends on the passage, as indicated by the two significant interaction terms in table 3.

**Fig. 2.**
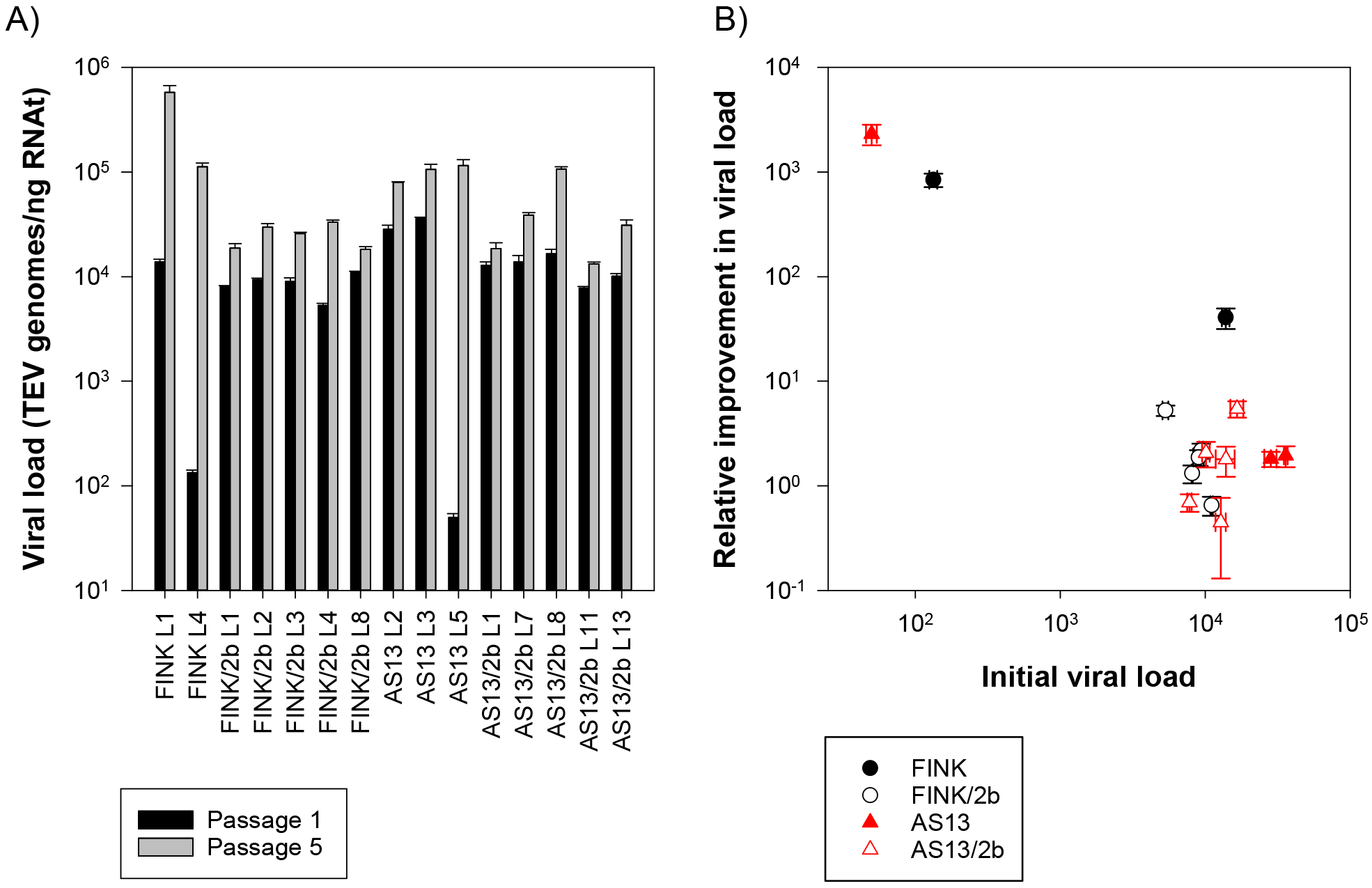
(A) Change in viral load (a proxy of within-host replicative fitness) observed between the first and the last evolution passages. (B) Negative correlation between viral load of the non-evolved genotypes and the magnitude of improvement at the end of the evolution experiment. In both panels, error bars represent ±1 SD.

**Table 3.**
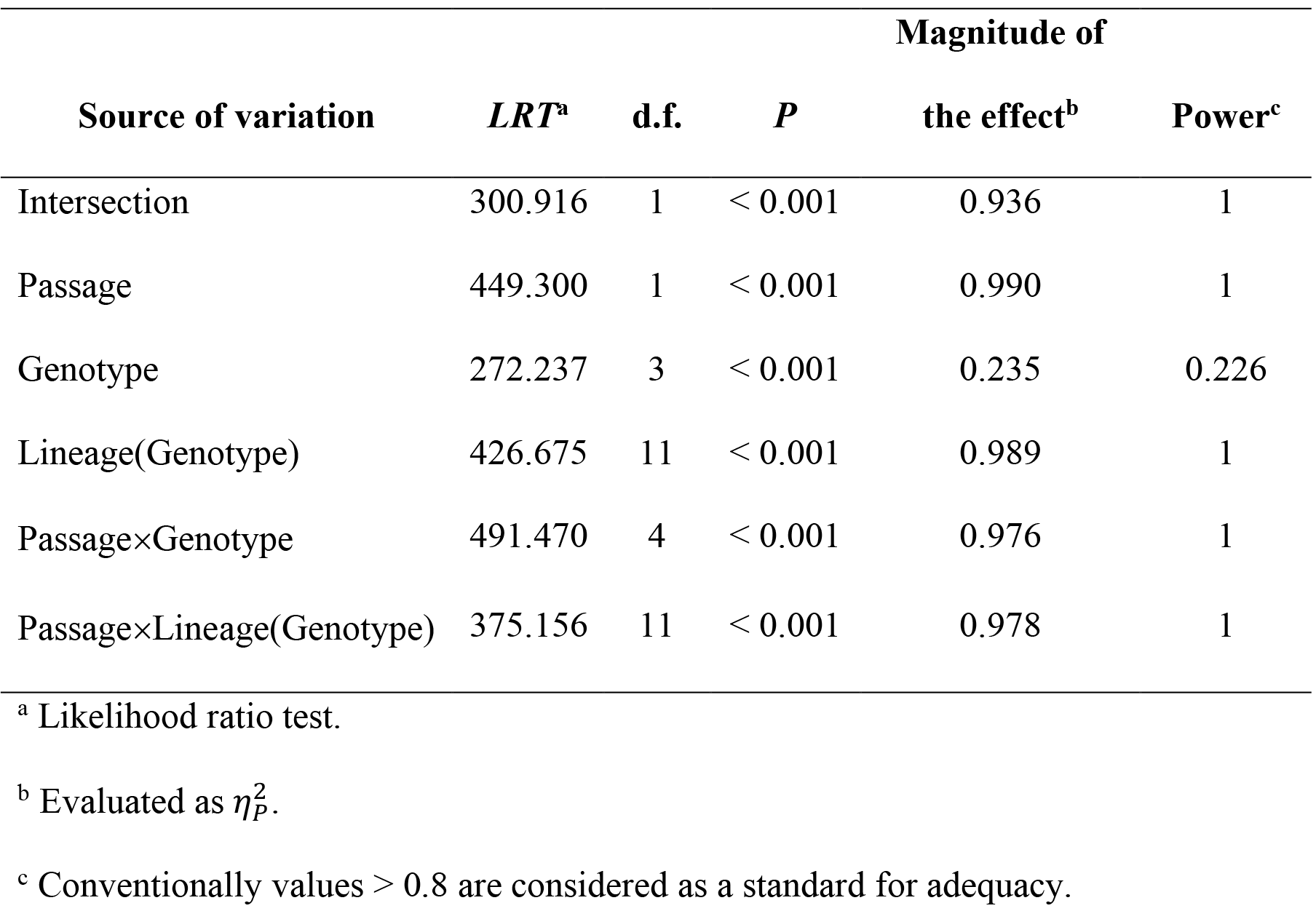
Generalized linear model testing for the effect of the different factors on viral load (within-host fitness). The fitted model assumes a Gamma distribution of errors and a logarithmic link function and is shown in equation 4.

Next, we sought to evaluate whether the observed changes in fitness were directly proportional to the magnitude of the perturbation induced by the corresponding mutations in HC-Pro RSS activity, as it would be expected from reversions to the wild-type sequence, or pervasively deviate from the linear relationship, as expected for the accumulation of compensatory mutations at other sites in the HC-Pro (or, in principle, elsewhere in the genome) (Moore et al. 2000; Sanjuán et al. 2005). To test this hypothesis, we first computed the relative change in viral load as *R*_*L(G)*_ = *V*_*L(G)*_ (5)/*V*_*L(G)*_ (1) − 1, where *V*_*L(G)*_(*t*) is the viral load estimated for lineage *L* of genotype *G* at the indicated *t* passage number. As illustrated in fig. 2B, the lower the viral load measured at passage 1, the larger the relative gain in fitness (linear regression: *R*^2^ = 0.754, *F*_1,13_ = 39.890, *P* < 0.001), thus supporting the null hypothesis of fitness recoveries being proportional to the actual deleterious effect. Indeed, fitting a quadratic model to the data in fig. 2B does not significantly improve the goodness of fit (partial *F* test: *F*_1,12_ = 3.686, *P* = 0.079), thus further rejecting the alternative hypothesis of second-site compensatory mutations. This conclusion of a predominance of reversions over second-site compensatory mutations will be confirmed in the sections below by determining the sequence of *HC-Pro* in all lineages.

### Stability of the *2b* Insert

Willemsen et al. (2017) found that, among several other exogenous coding sequences inserted in the genome of TEV, *2b* was particularly stable and functionally able to compensate the severe fitness costs associated with mutations in fundamental residues of HC-Pro. Here, we have evolved engineered genotypes of TEV encoding *2b* in a situation in which its RSS activity would be required to complement the deleterious effect of mutations in the virus’ natural RSS HC-Pro. However, a possible outcome from our evolution experiment would be the evolved viruses to recover fitness by compensating or reverting mutations in *HC-Pro* and subsequently removing partial or totally the *2b*. To explore this possibility, and the evolutionary stability of *2b*, we have confirmed the presence of *2b* in all the corresponding evolved lineages by RT-PCR using a pair of primers specifically designed to amplify the region into which *2b* was inserted (see Material and Methods for details). Fig. 3A shows the results of this study in all lineages after passage 1 and fig. 3B the same but after the last passage 5. All evolved *2b*-carrying lineages have retained the insert (all have the expected amplicon size of 651 bp). Therefore, this experiment confirms the evolutionary stability of the *2b* inserted into the TEV genome and that the observed patterns of recovery of fitness and virulence did not rely in removing this exogenous genetic material but in point mutations.

**Fig. 3.**
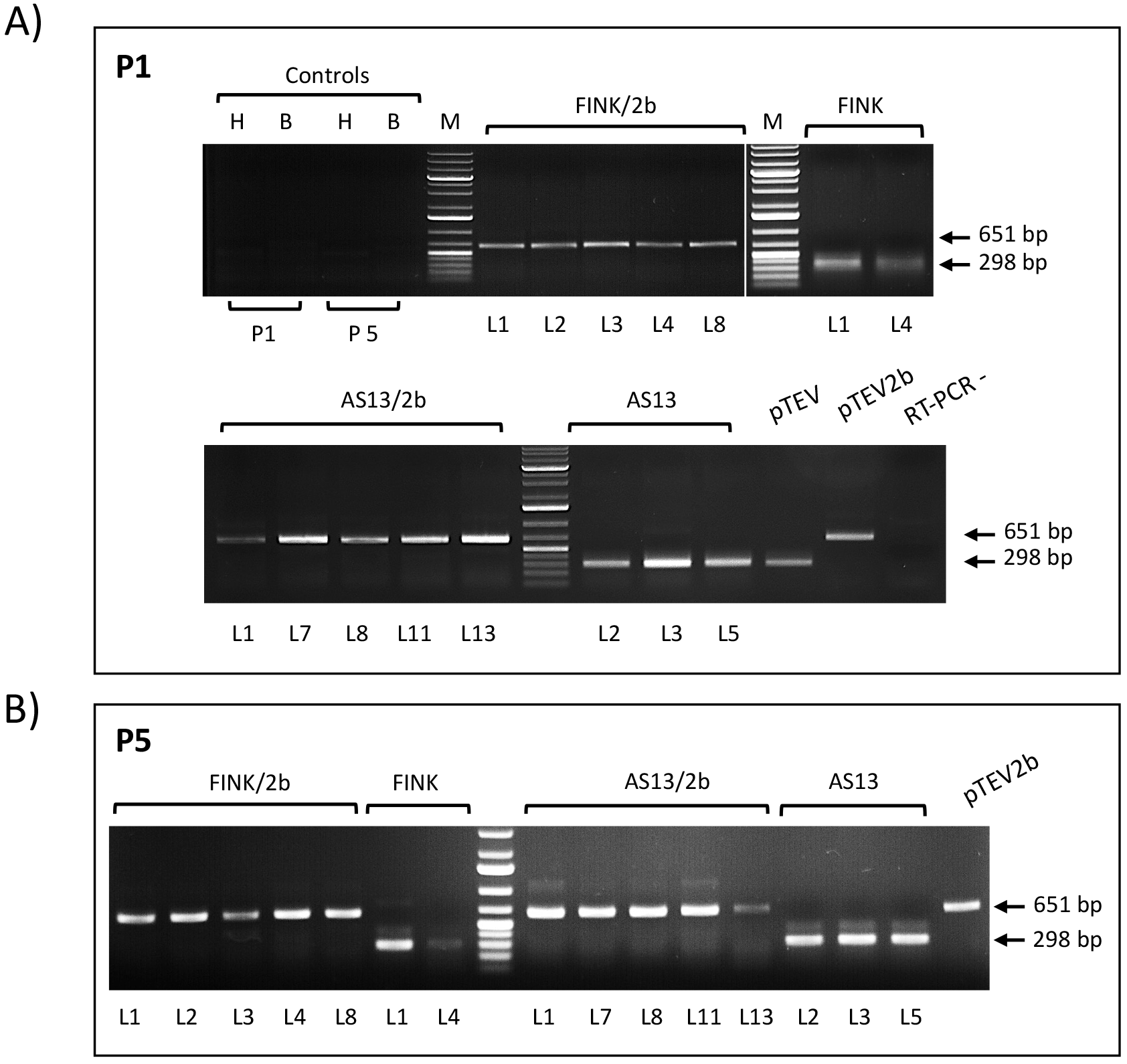
Confirmation of the presence of *2b* in the evolved viral lineages. Agarose gels (1%) with conventional RT-PCR products of the region encompassing the *2b* inserted into the TEV genome. Expected amplicon sizes are indicated on the right panels with arrows: 298 bp for wild-type TEV and 651 bp for TEV/2b. No deletions or reorganizations were observed in any lineage or passage. (A) Amplicons of all vims lineages from the first (P1) passage. (B) Amplicons from the fifth (P5) passage. Healthy (H), buffer-inoculated (B) plant extracts and water (RT-PCR-) were used as negative RT-PCR controls. PCR products obtained from the original full-length cDNA virus clones of TEV (pTEV) and TEV/2b (pTEV2b) are included as positive controls of insertion size. RT-PCR -: negative RT-PCR control.

Very interestingly, not a single mutation, not even in a polymorphic state has been observed in the *2b* gene (fig. 4C), thus supporting the high evolutionary stability of this exogenous gene into TEV genome.

**Fig. 4.**
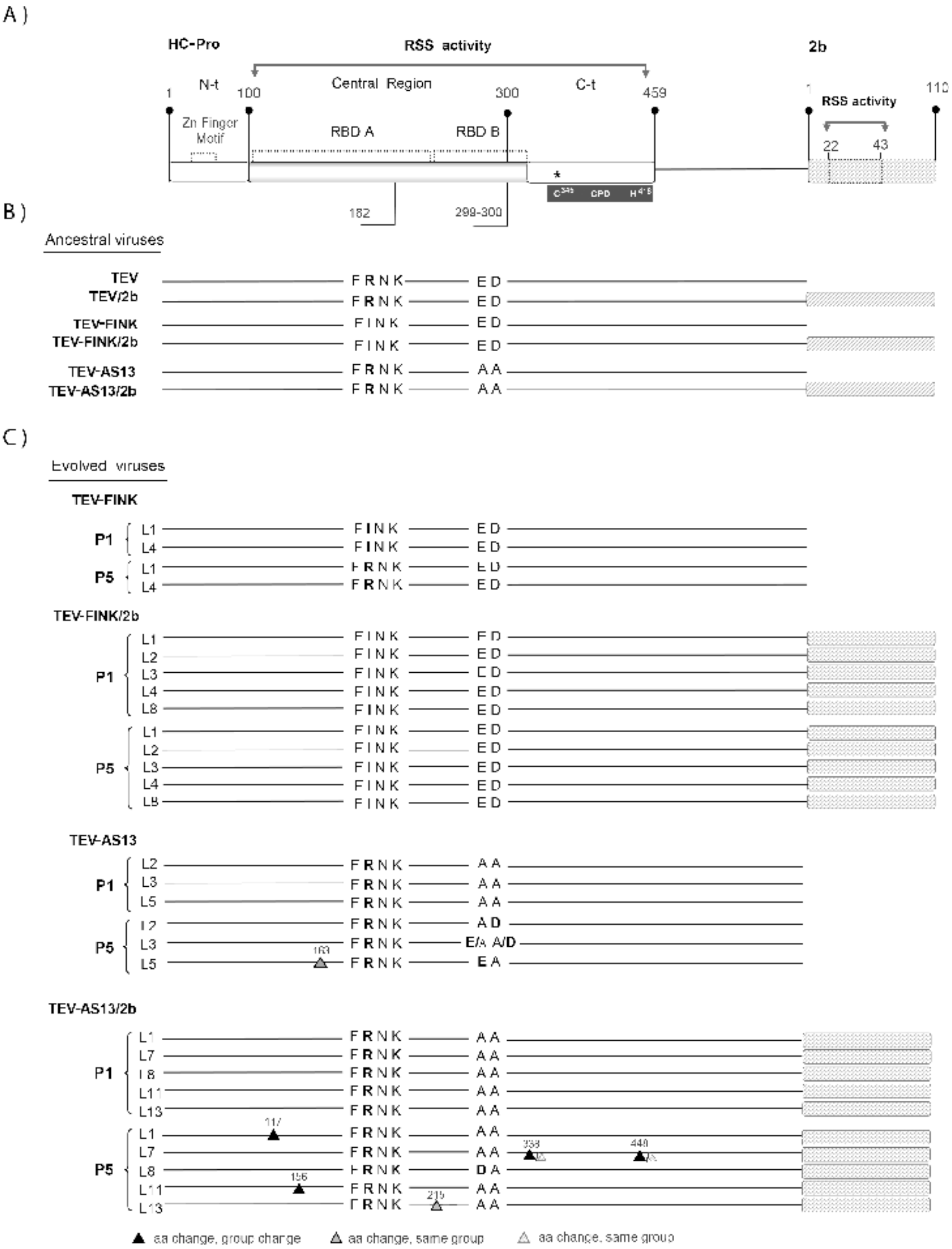
(A) Schematic representation of TEV HC-Pro and CMV 2b viral suppressors of RNA silencing proteins as present in the engineered TEV-FINK/2b and TEV-AS13/2b genomes. The main structural and functional region/domains and relevant motifs of HC-Pro and 2b are indicated. RSS: RNA silencing-suppression region. RBD: RNA-binding domains within the RSS. CPD: cysteine proteinase domain. Position of the TEV FRNK and ED amino acid motifs indicated with a horizontal line below the corresponding RBD regions. Position of the C345 and H418 dyad amino acids of the catalytic CPD in the Ct of the central region, are indicated by an asterisk. (B) Ancestral virus genotypes used in this work with the relevant amino acids and *2b* insertion marked by a gray box. (C) Same as in panel B but indicating the mutations found in the two genes in evolving viral lineages after one (P1) and five (P5) passages. The position and type of additional mutations found are marked with triangles. Large triangles denote amino acid replacements fixed in the consensus sequence (> 50%) while small ones represent minority viral subpopulations. For additional details see Supplementary Table S1.

### Molecular Changes Observed in HC-Pro

To characterize possible point mutations responsible for the observed phenotypic recoveries in presence and absence of genetic redundancy, we have obtained the consensus sequences of the *HC-Pro* and *2b* genes in all fifteen evolved lineages. Supplementary Table S1 shows all observed mutations. For guiding purposes, fig. 4A shows a schematic representation of the *HC-Pro* and *2b* genes, indicating the two HC-Pro RBD involved in RSS activity and the precise location of the two motifs mutagenized on each genotype (fig. 4B). Fig. 4C shows a schematic representation of the consensus sequences obtained for all fifteen evolved lineages at passages 1 and 5. The two TEV-FINK lineages had reverted the strong hyposuppressor mutant R183I to the wild-type arginine (fixed change), thus restoring the FRNK motif. In sharp contrast, none of the five TEV-FINK/2b shows such a reversion after passage 5, all retaining the mutated FINK sequence motif.

Regarding the three TEV-AS13 lineages, all had partially reverted the two deleterious mutations in a complex manner. Firstly, lineage L2 only reverted mutation D300A to the wild-type aspartic acid, being the change fixed. Secondly, lineage L5 only reverted mutation E299A to the wild-type glutamic acid residue, being also a fixed substitution. In addition, L5 also incorporates a fixed I163L (non-charged hydrophobic by non-charged hydrophobic) nonsynonymous mutation in the RBD A domain. Thirdly, lineage L3 shows polymorphisms in residues 299 and 300 of the protein alternating E299A/A299E and D300A/A300D. In both cases, the most frequent mutation corresponds to the wild-type allele (glutamic or aspartic acids, respectively), thought with the consensus sequencing approach here employed we cannot distinguish between having two or four haplotypes. Again, in sharp contrast, none of the TEV-AS13/2b lineages had reverted the knockdown alanine mutations into the wild-type amino acids. However, lineage L8 has incorporated a mutation to aspartic acid into residue 299, which is not a strict reversion, thought it brings back a negative charge into this motif. All other TEV-AS13/2b lineages contain additional nonsynonymous mutations elsewhere in the *HC-Pro* (fig. 5C). Lineage L1 has a C117Y (non-charged of moderate hydropathy by non-charged hydrophobic) fixed substitution in the RBD A domain. Lineage L11 has a A156T (non-charged hydrophobic by non-charged hydrophilic) fixed substitution in the RBD A domain. Lineage L13 has a A215V (non-charged hydrophobic by non-charged hydrophobic) fixed substitution in the RBD B. Interestingly, lineage L7 has two polymorphic amino acid substitutions, Y338C (non-charged hydrophobic by non-charged of moderate hydropathy) that is close to the C345 active site of the cysteine proteinase domain (CPD) (Oh and Carrington 1989; Kasschau et al. 1997), and G448D (non-charge hydrophobic by negatively charged hydrophilic) in the C-terminal region outside the RBD B domain (again, our consensus sequencing approach does not allow to conclude if two or four alleles are coexisting in the viral population).

### Association Between Virulence and Viral Accumulation

Although this section deviates from the main discussion about the role of genetic redundancy on compensatory evolution, we believe it is worth present and discussing, though in short, an interesting observation. A still controversial hypothesis brought forward to explain the evolution of parasite’s virulence assumes the existence of a positive association between virulence and within-host accumulation (Froissart et al. 2010). Increased accumulation will facilitate transmission at the cost of damaging the host. However, observations with plant viruses are highly variable regarding their support to this hypothesis. In some cases, a lack of positive association has been observed between accumulation and virulence (*e.g.*, Moury et al. 2001; Carrasco et al. 2007). In other cases, the predicted positive correlation was there (*e.g.*, Doumayrou et al. 2012) or found to be dependent on the particular combination of host- and virus-genotypes (*e.g.*, Pagán et al. 2007; Agudelo-Romero et al. 2008). Contributing to confusion, in some other cases even a negative correlation was found (*e.g.*, Rodríguez-Cerezo et al. 1991; Poulicard et al. 2010). Given that we have estimated virulence and within-host replication for a set of viral genotypes, it is possible to test this hypothesis in our particular pathosystem. A significant positive partial correlation, controlling for virus genotype, gave support to the hypothesis (*r*_*p*_ = 0.514, 27 d.f., *P* = 0.004).

## Discussion

The evolutionary consequences of genetic redundancy have been discussed at length. On the one hand, genetic redundancy may be the raw material for natural selection to evolve new functions (Nowak et al. 1997; Kondrashov et al. 2002) or to evolve moonlighting proteins with overlapping yet suboptimal activities (Lynch and Force 2000; Fares 2014; Espinosa-Cantú et al. 2015). On the other hand, by creating mutational robustness, genetic redundancy may contribute to pseudogenization and gene lost, specially under strong selection for genome reduction, as it may be the case for intracellular parasites (Mendoça et al. 2011), including viruses. In this study, we have explored the role of genetic redundancy for an essential viral function, the ability to suppress the RNA silencing defense of the host cells. After engineering TEV genomes containing deleterious mutations in their own RSS (a strong functional knockdown in the FRNK motif of RBD A or a weak one in the ED motif of RBD B; fig. 4B), we generated additional genomes expressing an exogenous RSS, the CMV *2b* gene. After evolving independent lineages from each one of these four genotypes, we evaluated virulence- and fitness-related traits and characterized the consensus sequence for *HC-Pro* and *2b* in each evolved lineage. We have observed that genetic redundancy affects the rate of compensatory evolution both at the phenotypic and at the genotypic level. At the phenotypic level, we have observed that, on average, genetically redundant viruses expressing the 2b RSS were more virulent and had higher fitness before starting evolution and experienced relatively smaller changes in these traits by the end of the evolution experiment (fig. 1B and fig. 2A). By contrast, non-redundant genotypes had very low virulence and fitness at the beginning of the evolution experiment and, by the last passage, had reached values similar to those of the redundant genotypes (fig. 1B and fig. 2A).

Even though our genotyping study was limited to population consensus sequences, we made a number of interesting observations (fig. 4C). First, the patterns of genomic evolution were completely different for genetically redundant and non-redundant genomes. While the strongly deleterious mutation FINK reverted to the wild-type motif FRNK in all the non-redundant evolved genomes, not a single reversion was observed in the redundant ones. Likewise, the mutations E299A and D300A partially reverted (to glutamic acid at position 299 or to aspartic acid at position 300 and it may have fully reverted in lineage L3), but never in the redundant genomes. Interestingly, the TEV-FINK/2b lineages did not show additional mutations anywhere else in the HC-Pro cistron, while 4 out of 5 TEV-AS13/2b lineages had fixed a number of nonsynonymous mutations affecting the RBD A (L1 and L11), RBD B (L13) or the C-terminal domain (2 mutations in L7). At this stage, the observation of additional mutations fixed along HC-Pro in the TEV-AS13/2b lineages is compatible with the two hypotheses exposed above. On the one hand, according to the hypothesis of genetic redundancy boosting the evolution of new functions, these extra mutations may be improving some of the additional functions performed by HC-Pro, after 2b taking the role as RSS (which will support the hypothesis of genetic redundancy allowing to generate or improve novel functions). In this way, stands out the two mutations observed in lineage L7 (fig. 4C) affecting the protease and movement domains, specially the first which renders a new cysteine residue (T338C) close to the C345 active site of the CPD (Oh and Carrington 1989; Kasschau et al. 1997). Whilst the second, a G448D, involves a replacement in the SGLE phosphoserine motif without affecting the active serine. In addition, mutations fixed in lineages L1, L11 and L13 also affect the movement domain (fig. 4C) (Plisson et al. 2003; Valli et al. 2018). On the other hand, according to the hypothesis of genetic redundancy buffering the effect of deleterious mutations and driving to pseudogenization and gene lost, the observed mutations may have a deleterious effect. However, under the pseudogenization hypothesis, we would expect to also observe fixation of synonymous mutations and, based on previous observations (reviewed in Willemsen et al. 2018), a number of in-phase deletions. Given that this is not the case, we believe our data, albeit circumstantially, are better supporting the first hypothesis. In any case, for both hypotheses, an acceleration in the rate of molecular evolution is expected in the genetically redundant lineages (Kondrashov et al. 2002), as we have observed for the TEV-AS13/2b ones. Clearly, site-directed mutagenesis experiments followed by biochemical and physiological characterization of the protease and movement activities of these mutants would help to distinguish between these two competing hypotheses.

Interestingly, we have also observed that the magnitude of the phenotypic improvement was proportional to the original fitness effect (fig. 2B): non-redundant genotypes (TEV-FINK and TEV-AS13) recovered proportionally more fitness than their corresponding genetically redundant counterparts (TEV-FINK/2b and TEV-AS13/2b, respectively). This proportionality suggests that most of the mutations involved in the virulence- and fitness-recovery process were reversions rather than second-site compensatory mutations (Moore et al. 2000; Sanjuán et al. 2005). Compensatory mutations are defined as those providing a disproportionally large fitness effect in the mutant genetic background compared to their effect in the wild-type background (*i.e.*, sign epistasis), coupled with retention of the original deleterious mutation. In this regard, our consensus sequencing results show a combination of effects: non-redundant genotypes have reverted mutations to the wild-type alleles, while redundant TEV-AS13/2b genotypes (but not TEV-FINK/2b) have fixed extra mutations that, as we just discussed in the previous paragraph, may be compensatory if they are improving the protease and/or movement activities.

An interesting question that pops up from our results is the clear difference in patterns of molecular evolution between the strong and weak hyposuppressor mutants. Lineages with the more deleterious mutation in the RBD A FRNK motif either revert to wild-type (TEV-FINK) or remained unchanged in presence of 2b (TEV-FINK/2b). By contrast, lineages with the weaker mutation in the RBD B ED motif were more variable in the observed outcome, as just discussed in the previous two paragraphs. Why is so? FRNK has been shown to be essential for the RSS activity of HC-Pro by directly binding to the siRNAs (Shiboleth et al. 2007; Kung et al. 2014). Therefore, mutations affecting this motif are completely abolishing the RSS activity (Shiboleth et al. 2007; Kung et al. 2014). In non-genetically redundant lineages, selection for reversion is maximally strong, thus almost deterministically ensuring the fixation of the required amino acid change. In the genetically redundant lineages, by contrast, RSS activity is over taken by 2b, relaxing the necessity of reverting isoleucine into arginine in the FRNK motif. By contrast, mutations in the ED motif retain a residual RSS activity (Torres-Barceló et al. 2008) and may be involved into other functions without fully removing them. This being the case, the nonredundant genotypes are under weaker selection than in the case of the FRNK motif and this is why we observe partial reversions and fixation of additional mutations. This observation is in good agreement with a previous compensatory evolution experiment performed with TEV-AS13 in which all evolved lineages fixed second-site compensatory mutations, but not reversions, (Torres-Barceló et al. 2010).

Another intriguing observation, aligned with the previous work by Willemsen et al. (2017), is the high genetic and evolutionary stability of the *2b* insertion. Remarkably, in both studies, the *2b* has been always retained without even fixing a single nucleotide substitution. What fitness benefit does *2b* provide to TEV? A first possibility would be that genetic redundancy increases mutational robustness. However, as recently shown by Willemsen et al. (2018), the mutational robustness of engineered TEV genomes expressing additional genes is inversely proportional to genome length and in the case of *2b* was not different than wild-type (see fig. 3 in Willemsen et al. 2018). Therefore, an increase in mutational robustness is not a likely benefit of expressing the 2b protein. A second possibility is that 2b facilitates TEV replication and/or cell-to-cell and systemic spread by interfering with the plant silencing suppression pathway in stages other than those affected by HC-Pro. HC-Pro sequesters sRNAs duplexes, relieving translational repression by the endonuclease AGO1, and inhibits the sRNA methylation activity of the methylase HEN1 (Ivanov et al. 2016; Valli et al. 2018). CMV 2b physically binds to AGO1 and provokes inhibition of the cleavage activity of the RISC machinery (Zhang et al. 2006) and also inhibits the production of RDR1-dependent viral siRNAs that confer systemic resistance by directing non-cell autonomous antiviral silencing (Guo and Ding 2002; Díaz-Pendón et al. 2007). Therefore, both RSSs proteins have overlapping functions but also differ in some aspects of their mechanisms of action, thus supporting the possibility of the *2b* insertion was selected under our serial passages regime because it increased virus accumulation beyond the level of HC-Pro by itself.

HC-Pro is a multifunctional protein composed by domains involved in overlapping functions (Plisson et al. 2003; Valli et al. 2018): mediator in vector transmission, RSS, cell-to-cell movement, and cysteine protease. It has been proposed that multifunctionality conflicts with optimization of every function (Guillaume and Otto 2012; Fares 2014) and therefore a tradeoff between all these functions is expected. Engineered genetic redundancy relaxes these evolutionary constraints and allows multifunctional proteins to optimize the functions that are not being provided *in cis*, in our case the RSS. The results we are reporting here for the TEV-AS13/2b lineages may represent an early step in this division of labor and specialization molecular evolutionary pathway. Longer evolution experiments and additional characterization of the adaptive value of fixed mutations will seed light into this interesting question.

## Acknowledgments

We thank Paula Agudo for excellent technical assistance. This work was supported by Spain’s Agencia Estatal de Investigación - FEDER grant BFU2015-65037-P to S.F.E. and by a fellowship from the Dominican Republic’s Ministerio de Educación Superior, Ciencia y Tecnología to S.M.R. The funders had no role in study design, data collection and analysis, decision to publish, or preparation of the manuscript.

